# An LLM-driven pipeline for proteomics-based detection and structural modeling of post-translational modifications

**DOI:** 10.64898/2026.05.01.722279

**Authors:** August George, Daniel Mejia-Rodriguez, Xiaolu Li, Paul Rigor, Margaret S. Cheung, Aivett Bilbao

## Abstract

Post-translational modifications (PTMs) on proteins dynamically regulate their functions and subsequently cellular physiology. Significant advances have been made in their detection and modeling: mass spectrometry-based proteomics has become the cornerstone for PTM detection in complex samples, while emerging structure-prediction frameworks enable modeling of PTM-dependent conformational changes. However, the biological significance of many PTMs remains largely unexplored, in part because integrated pipelines that bridge PTM detection with structural modeling remain limited. We present a generative AI-driven pipeline that integrates PTM detection with structural modeling of their effects on protein dynamics and interactions. The pipeline comprises two complementary tools: *PTMdiscoverer* and *PTM-Psi*. First, *PTMdiscoverer* leverages large language models to identify, annotate, and interpret candidate PTMs from open-search proteomics results, addressing limitations of conventional proteomics tools. Next, *PTM-Psi* models the structural, functional, and dynamic consequences of these spatially aware modifications on protein dynamics. These two components bridge PTM discovery with mechanistic interpretation at the structural level. We demonstrate our pipeline by using cyanobacterial proteomics data to study potential molecular mechanisms of redox-regulated “dark complex” formation in carbon metabolism, advancing our ability to interpret PTM-mediated regulation in microbial systems.

## INTRODUCTION

Protein function in microorganisms is commonly modulated by post-translational modifications (PTMs), which play vital roles in various cellular processes such as protein synthesis and turnover, metabolism, and cell cycle ^1^. Specifically, in cyanobacteria, PTMs help cells balance carbon flux across day-night cycles^2^. Phosphorylation and thiol-based PTMs link environmental light cues to light-dependent carbon metabolism and redox homeostasis. These effects are evident during diurnal growth and become acute under light-dark transitions, where misregulation leads to redox crisis and lethality^3,4^.

Bottom-up proteomics by liquid chromatography-mass spectrometry (MS) remains the predominant approach for PTM discovery in complex samples^5–7^, yet conventional data-analysis workflows still struggle with the breadth of PTM chemistry, and their outputs require labor-intensive expert curation. Rather than manual data curation, generative artificial intelligence (AI), specifically large language models (LLMs) based on transformer architectures^18^, significantly accelerate the integration of low-level numerical signals from MS with high-level contextual understanding, which enables a more holistic approach for deeper insights into biological processes^8^. Furthermore, existing proteomics software packages rarely integrate structural interpretation, leaving the impact of PTMs on protein structure and dynamics largely unexamined and limiting our mechanistic understanding of PTM-mediated regulatory processes.

PTM-Psi, a python package to facilitate the computational investigation of post-translational modification on protein structures and their impacts on dynamics and functions, has previously been developed and validated as a structural modeling framework for evaluating PTM-induced conformational and dynamic effects^9^, and has recently been scaled for high-throughput with cloud computing^10^. However, its application has been limited by the lack of automated, biologically contextualized PTM prioritization from proteomics data.

Here we introduce an LLM-driven pipeline addressing these limitations by integrating *PTMdiscoverer*^11^ for proteomics-based PTM detection, and *PTM-Psi*^9^ for structural modeling of PTM impacts. PTMdiscoverer leverages large language models to detect, filter, and annotate candidate PTMs from open-search proteomics data, and produces structured, residue-resolved PTM annotations that can be formatted as inputs for structural modeling with PTM-Psi. A case study of the glyceraldehyde-3-phosphate dehydrogenase (GAPDH)/CP12/phosphoribulokinase (PRK) protein megacomplex in cyanobacteria, a so-called “dark complex”^12^, is presented to generate mechanistic hypotheses for carbon-metabolism regulation in *Synechococcus elongatus* under light disturbance. We focus on PTMs implicated in circadian-controlled and light-responsive pathways, illustrating how complementary detection and modeling tools could connect PTM events to enzyme conformational states. This framework aligns with emerging structural insights into cyanobacterial redox control^13^ and provides a conceptual workflow generalizable to PTM regulation across bacterial systems.

## RESULTS AND DISCUSSION

To illustrate our pipeline, we re-analyzed a subset of MS-based bottom-up proteomics data acquired to investigate the role of PTMs in the regulation of carbon metabolism in *Synechococcus elongatus* under light disturbance^14^. Raw measurement data are openly accessible at the Mass Spectrometry Interactive Virtual Environment (MassIVE) repository under accession MSV000096184. Three proteins of the GAPDH/CP12/PRK “dark complex” were targeted: GAP2 (Q9R6W2), CP12 (Q6BBK3), and PRK (Q9LBV7).

The reference proteome of *S. elongatus* (UniProtKB taxonomy ID 1140, 2025/02/25) plus unreviewed sequences for these proteins were used.

The workflow is summarized in Figure 1. MSFragger (v.4.1) was used to identify proteins at 1% false discovery rate (FDR) via traditional search and to identify delta masses via open search^15,16^. Delta masses identified through open search were filtered through a multi-stage quality and localization cascade (see Methods), yielding 92 (CP12), 394 (PRK), and 171 (GAP2) candidate modification events. A zero-shot prompt incorporating domain knowledge, experimental context, and chain-of-thought reasoning instructions was constructed per protein (see Methods; Supplementary File 1). PTMdiscoverer performed batch LLM inference using the gpt-4o model via the OpenAI API to prioritize these candidates into a concise subset of experimentally supported and biologically plausible PTMs (Table 1).

**Table 1.** PTMs detected for downstream structural modeling to investigate carbon metabolism regulation of *S. elongatus* under light disturbance. Where specific residue positions are listed (e.g., CP12 Cys22, Cys31, Cys33, Cys64), these were unambiguously mapped to absolute protein coordinates using MSFragger localization data; positions reported as general regions indicate cases where peptide-to-protein mapping was ambiguous.

| Gene | AI-annotated PTM Name | $\Delta$ Mass (Da) | Representative Residues | AI-Annotated Functional Relevance |
| --- | --- | --- | --- | --- |
| GAP2 | Oxidation | 15.995 | Cys residues (e.g., catalytic/near-catalytic) | Redox-sensitive modulation of glycolytic/Calvin-cycle flux |
| GAP2 | NEM Alkylation | 57.029 | Cys residues | Experimental thiol blocking consistent with redox-state probing |
| CP12 | Oxidation | 15.994 | Cys22, Cys31, Cys33, Cys64 | Redox-dependent complex formation with GAPDH/PRK |
| CP12 | NEM Alkylation | 125.044 | Cys residues | Thiol alkylation consistent with experimental NEM blocking |
| PRK | Oxidation | 15.997 | Cys-containing regions | Possible oxidative tuning of PRK catalytic activity |
| PRK | NEM Alkylation | 57.023 | Cys residues | Experimental thiol labeling |

**Figure 1.**
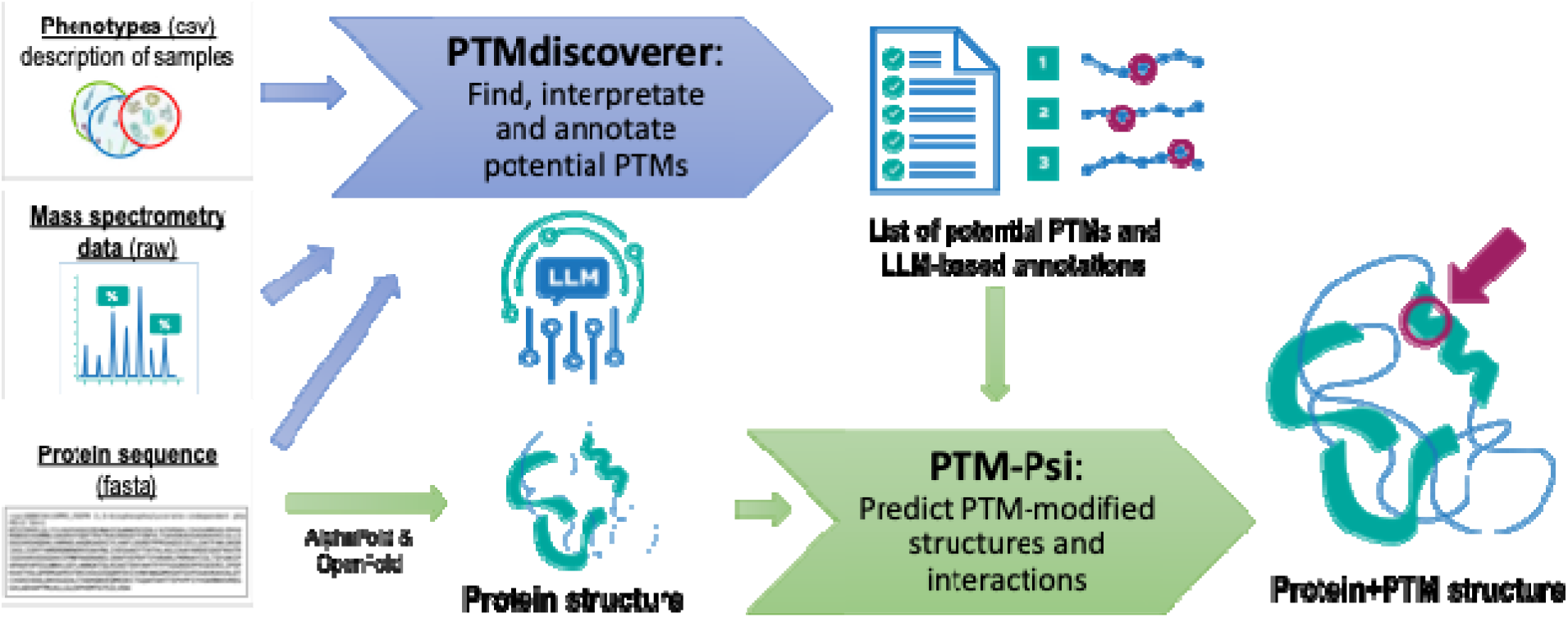
Conceptual two-stage workflow for proteomics-based PTM detection and structural modeling. <u>Top - PTMdiscoverer:</u> Raw proteomics MS data are first processed in open search mode to generate tables with peptide-spectrum matches and delta masses corresponding to tentative PTMs, which PTMdiscoverer then filters through a multi-stage quality and localization cascade. This enables recognition of both known and previously elusive modifications. Delta masses are evaluated by LLM inference to produce a prioritized list of biologically relevant PTM annotations, including residue positions, modification types, and hypothesized functional effects. <u>Bottom - PTM-Psi:</u> The protein sequence is used to predict a 3D protein structure. The PTM annotations from PTMdiscoverer and the predicted protein structure serve as inputs to model PTM-modified protein conformations, dynamics, and interactions. Arrows between tools indicate data flow.

Table 1 summarizes the results from PTMdiscoverer with literature-supported evidence. Multiple redox-associated modifications were identified across GAP2, CP12, and PRK, including oxidation events with delta masses of 15.994–15.997 Da. The +15.99 Da mass shift detected on cysteine residues across all three dark complex proteins is consistent with cysteine oxidation, likely sulfenylation (Cys-SH → Cys-SOH), aligning with the known redox regulation of this complex^12^. Interestingly, cysteine sulfenic acid is generally considered unstable and often seen as an intermediate step towards more stable oxidized forms^20-22^, such as those containing intra- or intermolecular disulfide bonds (cystine, glutathionylated cysteine), not identified by PTMdiscoverer. The detected NEM alkylation signatures (e.g., 57.029 Da and 125.044 Da), which reflect sample preparation rather than biological modifications, were consistent with thiol labeling strategies used to probe cysteine redox states. The predominance of cysteine-centered modifications aligns with the redox-regulated Calvin-cycle control mechanisms reported under light disturbance conditions^14^. These prioritized PTMs were consistent with previously reported results from structural modeling using PTM-Psi^10^, supporting their suitability for downstream perturbation studies.

Several limitations should be noted. First, LLM outputs are non-deterministic, meaning identical inputs may produce different annotations across runs. While systematic benchmarking was not performed in this study, PTMdiscoverer includes built-in multi-run consensus analysis to quantify this variability. Second, the tool currently depends on a commercial API (OpenAI gpt-4o) but supports configurable model and endpoint flags, enabling use with open-weight alternatives. Finally, this study examined three proteins from a single organism, and broader validation is needed to assess generalizability across diverse proteomes. Relative to tools such as PTM-Shepherd^16^ that statistically summarize delta-mass distributions, PTMdiscoverer adds LLM-driven biological context evaluation and functional annotation tailored to specific experimental conditions.

Beyond the case study presented above, PTMdiscoverer is designed for broad reuse. PTMdiscoverer is deployed as both a standalone command-line tool and an MCP-compatible (Model Context Protocol) server exposing six tools for study validation, protein listing, delta-mass extraction, LLM inference, result retrieval, and deterministic analysis. A web application built with Streamlit and the ADEPT^17^ (Agentic Discovery and Exploration Platform for Tools) framework provides an interactive interface in which a LangChain-based agent orchestrates PTMdiscoverer alongside complementary MCP tools for web search, sequence analysis, chemical database queries (PubChem, UniProt), and document retrieval-augmented generation (RAG). The application is containerized with Docker for reproducible deployment. Future work will deploy an end-to-end and automated integration with PTM-Psi for structural perturbation analysis and systematic benchmarking across diverse proteomes and language models.

Our pipeline aims to democratize proteomics data interpretation and structural modeling of PTMs, enabling the discovery of how environmental changes shape microbial function through PTMs, and accelerating insights into causal links between stimuli and biological processes across microbial systems. This work highlights a scalable and generalizable framework that lowers barriers to PTM analysis, demonstrates the potential of AI-assisted reasoning in proteomics, and provides a foundation for future integrative studies linking molecular modifications to protein structure, dynamics, and cellular function.

## METHODS

### Delta-mass extraction and filtering

PSM tables (psm.tsv) generated by MSFragger open search (v4.1) were processed per protein across all MS runs in the study. Rows were retained requiring zero missed cleavages, PSM probability > 0.9, and delta mass > 10.0 Da. Localization confidence required a single best position, best-position ion score ≥ second-best, and best-position score strictly greater than second-best. Delta masses were rounded to three decimal places and deduplicated per peptide, charge state, delta mass, best position, and MS run.

### LLM-based PTM annotation

A structured zero-shot prompt was constructed per protein by prepending a base prompt (Supplementary File 1) to the filtered delta-mass table. The prompt provides experimental context (TMT labeling, NEM thiol blocking, *S. elongatus* circadian conditions) and instructs the model to perform sequential analytical tasks including delta-mass-to-PTM mapping (±0.05 Da tolerance, 14-type controlled vocabulary), functional impact analysis, and generation of a validated JSON summary. Inference used the OpenAI API with the gpt-4o model. Output JSON was parsed, protein positions were augmented using MSFragger positional data when unambiguous, and records were validated for required fields.

### Software

PTMdiscoverer is a Python package providing a CLI with subcommands for study validation, protein listing, delta-mass extraction, inference, result inspection, and multi-run analysis, as well as an MCP-compatible server exposing six tools for agentic platform integration. The software is available at https://github.com/pnnl/PTMdiscoverer.

## SUPPLEMENTARY MATERIALS

Supplementary File 1: Base prompt template for LLM-based PTM annotation.

## AUTHOR CONTRIBUTIONS

August George: software development, prompt improvements, investigation, figure refinements, AI models, manuscript writing; Daniel Mejia-Rodriguez: investigation, validation; Xiaolu Li: LC-MS proteomics experiments; Paul Rigor: coding, AI models, and cloud support; Margaret S. Cheung: conceptualization, research design, manuscript editing, funding acquisition; Aivett Bilbao: conceptualization and coding of PTMdiscoverer, investigation, validation, figure, manuscript writing; All authors discussed the results, contributed to the manuscript, and approved the final version.

## ACKNOWLEDGMENTS

The authors thank Derek Munson for improving the figure. This research was funded by the Generative AI for Science, Energy, and Security Science & Technology Investment under the Laboratory Directed Research and Development Program at Pacific Northwest National Laboratory (PNNL), a multiprogram national laboratory operated by Battelle for the U.S. Department of Energy; Portions of this work were supported by the Center for AI at PNNL, the NW-BRaVE for Biopreparedness project funded by the U. S. Department of Energy (DOE), Office of Science, Biological and Environmental Research program, under FWP 81832, and performed at the Environmental Molecular Sciences Laboratory (EMSL), a DOE Office of Science User Facility sponsored by the Biological and Environmental Research program under Contract No. DE-AC05-76RL01830.

## AI DISCLOSURE

Generative AI tools were used in two capacities in this work: (1) as a core component of the pipeline, where large language models perform PTM annotation from proteomics data as described in the Methods, and (2) to assist with manuscript drafting, editing, and software development. All AI-assisted outputs were reviewed and approved by the authors, who assume full responsibility for this publication.

