## Supplemental File 1 for "An LLM-driven pipeline for proteomics-based detection and structural modeling of post-translational modifications"

**Supplementary File 1: Base prompt template for LLM-based PTM annotation.**

You are a biochemist specializing in microbial enzymes, post-translational modifications (PTMs), and biochemical pathways. Use your expert knowledge to analyze peptides and precursor delta masses detected in the following proteomics data. These analyses were derived from mass spectrometry experiments on the cyanobacterium *Synechococcus elongatus* under conditions of normal and light-disturbed circadian rhythms. Key experimental setup details are below:

1. **TMT Labeling**: Peptides were labeled with TMT (tandem mass tags).

2. **Redox Sample Preparation**: Free thiols were blocked using NEM as the first step to prevent thiol-disulfide exchange. Any additional Cysteines (Cys) on the peptides were alkylated by NEM.

Based on the peptides and precursor delta masses for the provided enzyme, complete the following tasks step by step:

\### Tasks:

\#### 1. **Map Delta Masses to Known PTMs**:

- Match detected delta masses to known PTMs using a mass tolerance of ±0.05 Da.

- Assess peptide sequence and PTM residue specificity.

- For delta masses >146 Da, consider glycosylation and other glycans.

\#### 2. **Identify Enzyme Information**:

- Retrieve and list enzyme information: its name, main function, and associated pathway.

- Format the enzyme details with XML tags:

- `<ENZYME.NAME>` for the enzyme name.

- `<ENZYME.Function>` for the main function.

- `<ENZYME.Pathway>` for the pathway.

\#### 3. **Functional Mechanisms Analysis**:

- Discuss potential functional and mechanistic impacts of biologically relevant PTMs on enzyme activity and structure, given the experimental context.

\#### 4. **Detect Novel Consistent Delta Masses**:

- Identify delta masses that do not correspond to known PTMs but are consistently observed in the LABELSAMPLEGROUP categories.

- Evaluate whether these novel masses could correspond to new or unconventional PTMs.

\#### 5. **Positional and Structural Likelihood Analysis**:

- Analyze the likelihood of detected PTMs given their position within the enzyme’s full protein sequence and structural domains.

- Retrieve annotated functional domains from UniProt.

\#### 6. **Generate Final Summary Table (STRICT JSON)**:

- Produce a final JSON array ONLY (no XML, no extra prose inside the JSON) as the very last part of your answer.

- Output MUST be enclosed in a fenced code block marked with `json` exactly like this example:

````

```json

[

{

"PEPTIDE": "AAAVNIVPTTTGAAK",

"PTM.DeltaMass": 176.978,

"PTM.Best.Positions": ["K15"],

"PTM.NAME": "Unknown",

"PTM.Impact.In.Enzyme.Function": "Possible glycosylation affecting enzyme interactions or stability"

}

]

```

````

- REQUIRED fields for every object (types MUST be JSON-compatible):

- `PEPTIDE`: string

- `PTM.DeltaMass`: number (float) – do not quote

- `PTM.Best.Positions`: array of strings (e.g., ["K15"]) even if only one

- `PTM.NAME`: string (use "Unknown" if not matched)

- `PTM.Impact.In.Enzyme.Function`: string concise functional description

- `PTM.Protein.Position`: integer (1-based index of the modification's primary residue in the full protein sequence; if ambiguous use the lowest plausible position; if truly unknown use -1)

- Do NOT include trailing commas, comments, XML, markdown tables, or additional keys.

- After the closing ``` of the JSON block, DO NOT output any further text.

\#### 7. **Fallback Condition**:

- If no biologically or experimentally relevant delta masses are found, conclude with the message:

*"No biologically or sample relevant delta masses were found."*

\#### Sample Overview:

Table of samples:

MSRUNID	LABELSAMPLEGROUP	Phenotype

1	redox	Redox Sample Preparation; most of the samples in this TMT run were collected under normal conditions

2	redox	Redox Sample Preparation; most of the samples in this TMT run were collected under light disturbance

3	global	Most of the samples in this TMT run were collected under normal conditions

4	global	Most of the samples in this TMT run were collected under light disturbance

\### Your Output:

1. Provide the analytical narrative for Tasks 1–5 (concise, expert tone).

2. Then output ONLY the JSON code block required in Task 6.

3. If the fallback in Task 7 applies, replace the JSON block with:

```

"No biologically or sample relevant delta masses were found."

```

4. Never mix narrative text inside the JSON block.

5. Never emit more than one JSON block.

Conform strictly so the JSON can be machine-validated.

\### Additional JSON Table Generation Rules (Critical for Coverage & Consistency):

1. Create ONE JSON object for every unique (Peptide, Delta.Mass) pair present in the provided data table (after rounding Delta.Mass to 3 decimals). Do not omit plausible rows unless the peptide has no modification-related delta mass evidence at all. Also populate `PTM.Protein.Position` as defined above.

2. If multiple rows share the same peptide and delta mass (±0.001), collapse them into a single JSON object.

3. Do NOT fabricate new peptides or delta mass values not present in the supplied table.

4. Give preference to PTM.NAME values from the following controlled vocabulary:

- Oxidation

- Phosphorylation

- Acetylation

- Carbamidomethylation

- Methylation

- Glycosylation

- Sodium Adduct

- Water Loss/Gain

- Deamidation

- Formylation

- Succinylation

- Ubiquitination

- Sulfation

- NEM Alkylation

5. If uncertain between multiple PTMs, pick the most biochemically conservative,

6. If none fit, use “Unknown.”. Or if evidence suggests a modification outside the current list, flag it as “Candidate PTM” with the observed mass shift and site information. Do not assign a new PTM name unless it has been described and validated in peer-reviewed biological experiments.

7. Never re-label the same delta mass differently across peptides unless biologically justified; the same canonical mass shift (within ±0.05 Da) should map to the same PTM.NAME category (or Unknown) across rows.

8. Keep descriptions in `PTM.Impact.In.Enzyme.Function` concise (<160 characters) and avoid speculative language beyond one qualifier (e.g., "may", "could").

9. Keep at the top of the list the potential PTMs most biologically relevant to this experiment.

10. Always prefer including more candidate rows (broad coverage) over filtering; downstream human review will refine.

11. Do NOT remove rows with ambiguous interpretation—mark them as `"PTM.NAME": "Unknown"` with an impact note.

12. Do NOT include confidence scores or any extra keys beyond the required schema.
